# Elevated nest temperature has opposing effects on host species infested with parasitic nest flies

**DOI:** 10.1101/2021.05.07.440146

**Authors:** Lauren Albert, Samantha Rumschlag, Alexandra Parker, Grace Vaziri, Sarah A. Knutie

**Author notes:** Department of Biology, Indiana University, Bloomington, IN USA. We demonstrate elevated temperatures have contrasting indirect effects on a parasite infesting two hosts. Our study provides novel implications of altered host defense to parasitism and understanding of the net-effects of climate change on host-parasite interactions. Author contributions: LA and SAK conceived the ideas and designed methodology; LA, AP, SAK collected data; SR and GV analyzed the data; LA, SAK, GV, SR contributed to the writing of the manuscript. All authors gave final approval of the manuscript.

## Abstract

Environmental factors, such as elevated temperature, can have varying effects on hosts and their parasites, which can have consequences for disease outcomes. The individual direct effects of temperature must be disentangled to determine the net-effect in host-parasite relationships, yet few studies have determined the net-effects in a multi-host system. To address this gap, we experimentally manipulated temperature and parasite presence in the nests of two host species infested by parasitic blowflies (*Protocalliphora sialia*). We conducted a factorial experiment by increasing temperature (or not) and removing all parasites (or not) in the nests of eastern bluebirds (*Sialia sialis*) and tree swallows (*Tachycineta bicolor*). We then measured nestling morphometrics, blood loss, and survival and quantified parasite abundance. We predicted that if temperature had a direct effect on parasite fitness, then elevated temperature would cause similar directional effects on parasite abundance across host species. If temperature had a direct effect on hosts, and therefore an indirect effect on the parasite, parasite abundance would differ across host species. Heated swallow nests had fewer parasites compared to non-heated nests. In contrast, heated bluebird nests had more parasites compared to non-heated nests. The results of our study demonstrate that elevated temperature can have differential effects on host species, which can impact infestation susceptibility. Furthermore, changing climates could have complex net-effects on parasite fitness and host health across multi-host-parasite interactions.

## 1 INTRODUCTION

Climate change can alter species interactions, including host-parasite relationships (Martinez & Merino 2011, Brooks & Hoberg 2007, Scharsack et al. 2015). Hosts and their parasites have evolved under specific abiotic and biotic conditions and a disruption to these conditions could have implications for the fitness of both the host and parasite (Musgrave et. al 2019, Wolinska & King 2009, Penczykowski, Laine, & Koskella 2015). Elevated temperature is an especially important factor in the effects of climate change on living organisms and their interactions (Møller et al. 2014). For example, because hosts and parasites have different optimal thermal breadths, a shift in temperature may concurrently affect the host and parasite differently (Paull, LaFonte, & Johnson 2012, Studer, Thieltges, & Poulin 2010). When individual effects of temperature on the host and parasite are considered together, these net-effects could alter the dynamics of how, or which, host-parasite interactions continue to exist in a changed environment.

Temperature can have direct and indirect effects on free-living parasite fitness, transmission rates, and exploitation of hosts (Martinez & Merino 2011, Bush et al. 2001, Hernandez, Poole, & Cattadori 2012, Amat-Valero, Calero-Torralbo & Valera 2013). For example, developmental times of Arctic nematodes *(Osteragia gruehneri)* slow as ambient air temperatures increase beyond a thermal maximum of 30°C, thereby lowering the density of the infectious stage (Hoar et al. 2012). Additionally, transmission rates can be negatively influenced by changing temperatures through an influence on the behavior or activity of free-living stages (Koprivinkar et al. 2010). Alternatively, parasites can be indirectly affected by temperatures through the positive and negative effects of temperature on hosts (Poulin 1998). Temperature can have contrasting effects on host species depending on their traits (Ferris & Best 2018, Ward, Kim, & Harvell 2007, Franke et al. 2017). Some host species may benefit from elevated temperatures, such as in the abalone (*Haliotis rubra*) whose antiviral activity and total cell counts in the hemolymph increased with temperature (Dang, Speck, & Benkendorff 2012). However, other host species experience detrimental effects to their fitness in response to elevated temperatures (Salaberria et al. 2014). For example, when temperatures reach the critical thermal maximum for amphibian hosts, they are more negatively affected by fungal pathogens (Greenspan et al. 2017, Cohen et al. 2017). The way in which hosts are influenced by temperature may concurrently affect their response to parasitism, creating the potential for complex effects in host-parasite interactions that could vary across host species.

Disentangling the causal effect of elevated temperature on multi-host-parasite interactions requires an experimental manipulation of temperature and the presence of generalist parasites across different host species. An ideal system to address this idea is with parasitic blowflies (*Protocalliphora sialia*) and their avian host species, tree swallows (*Tachycineta bicolor*) and eastern bluebirds (*Sialia sialis*). These host species are secondary cavity nesters that readily build nests in artificial cavities, such as nest boxes. Adult flies are non-parasitic but lay their eggs in bird nests once nestlings hatch. Flies develop through three larval instar stages while they are feeding non-subcutaneously on the blood and other secretions of nestling birds. Most studies have found no detectable, lethal effects of blowflies on nestling survival, while other studies have found sublethal effects of blowflies, including increased blood loss and decreased nestling growth (Grab et al. 2019). Although previous studies have demonstrated an influence of temperature on parasite abundance (Bennet & Whitworth 1991, Dawson et al. 2005), it is unclear whether there is a direct effect on the parasite or indirect effect via the host.

The goal of our study was to determine whether the effect of elevated temperature on parasite abundance is direct or indirect across hosts and whether an interaction between parasitism and nest temperature affects nestling health. We experimentally elevated nest temperature (hereon, heated) or not (hereon, non-heated) and manipulated blowfly presence by removing all blowfly parasites (hereon, non-parasitized) or allowing for natural parasitism (hereon, parasitized). We then quantified total parasite abundance and calculated parasite density (number of parasites per gram of nestling) to control for clutch size and body mass differences between host species. When nestlings were 10 days old, we measured nestling morphometrics (body mass, tarsus length, and first primary feather length). We also measured nestling hemoglobin levels, which are a more sensitive measure of the effect of parasitism on nestling health as a proxy of blood loss (O’Brien, Morrison & Johnson, 2001). Fledging success was also quantified for each nest as a measure of nestling survival.

We hypothesized that if nest temperature influences parasite fitness directly, then parasite abundance would change similarly across host species in response to elevated temperature (Figure 1A). Based on past studies, elevated temperatures could directly, negatively affect parasite fitness (Castaño-Vásquez et al. 2018). Alternatively, elevated temperatures could affect nestlings, which could have downstream, indirect effects on parasite fitness. Past work suggests that bluebirds and swallows respond to elevated temperatures differently, with heat decreasing bluebird fitness while increasing swallow fitness (Cohen et al. 2020, Sykes 2020, Dawson, Lawrie, O’Brien 2005). Therefore, we predicted that if elevated temperatures indirectly affect parasite fitness through the host, parasite abundance would differ across host species in response to temperature (Figure 1B, C).

**Fig 1.**
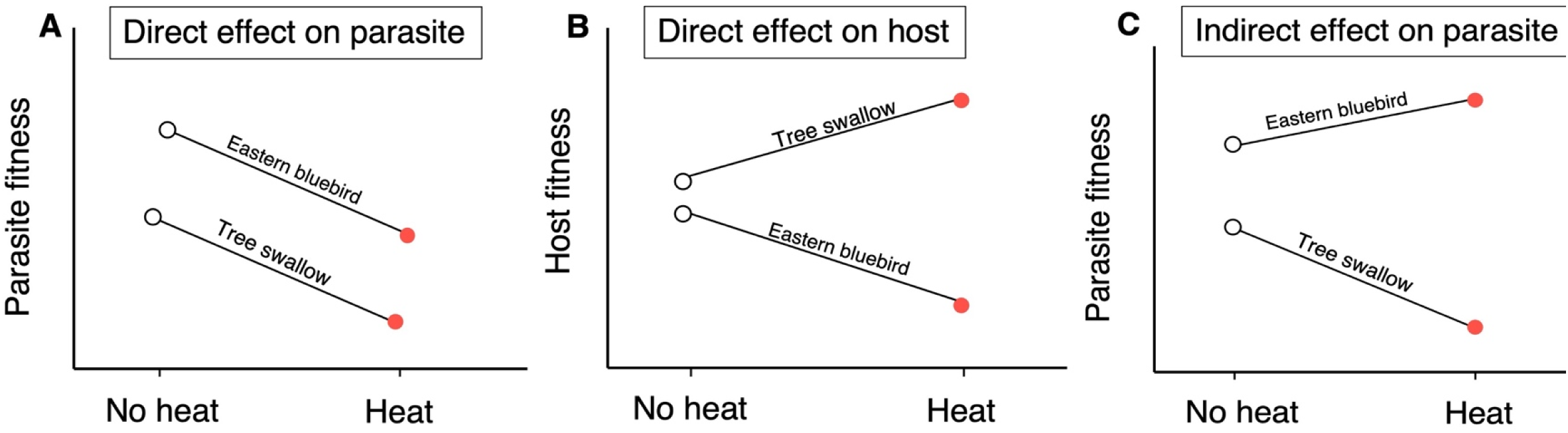
Conceptual figure of our predictions. A) A direct effect of heat on the parasite would cause similar directional effects of parasite fitness in the nests of both species. B) A direct effect of heat on host fitness could result in elevated temperatures negatively affecting bluebird fitness and positively affecting tree swallow fitness. C) An indirect effect of heat on the parasite would cause different directional responses of parasite fitness.

## 2 MATERIAL AND METHODS

### 2.1 Study system

Our study was conducted from May to August 2018 in northern Minnesota, USA near the University of Minnesota Itasca Biological Station and Laboratories (47°13’33”N, -95°11’42”W) in Clearwater and Hubbard Counties. In 2018, we observed approximately 170 nest boxes that are established haphazardly on the properties of landowners and within Itasca State Park. Nest boxes were mounted on metal poles or telephone poles and were either Peterson boxes or rectangular boxes, which vary in dimensions (Peterson boxes, bluebirds: n = 19, s wallows: n = 11; rectangular boxes, bluebirds: n = 19, swallows: n = 24).

Tree swallows and eastern bluebirds are abundant in the area and each nest readily in these artificial cavities. Tree swallows are long-distance migrants, returning to northern United States to brood from May-July in Minnesota (mean hatch date: 14 June 2018) (Sockman & Courter 2018). Tree swallow clutch size ranges from one to nine eggs, which are incubated for 13-14 days (Grab et al. 2019). Eastern bluebirds are short-distance, and sometimes partial migrants, returning to the eastern United States to breed from May-August (mean hatch date: 20 June 2019) (Sockman & Courter 2018). Bluebird clutch size ranges from one to seven eggs, which are incubated for 13-14 days (Grab et al. 2019). Minnesota tree swallow and bluebird nestlings spend an average of 20 and 18.8 days, respectively, in the nest where they are parasitized exclusively by blowflies (Grab et al. 2019, Gowaty & Plissner, 2015).

### 2.2 Experimental manipulation of parasites and temperature

Nest boxes were checked once a week for evidence of nesting activity (i.e., nest construction). Once eggs were found, clutch initiation date was determined by backdating one egg per day from the first record of eggs in the nest, and then nests were checked every other day until the first egg hatched. During the nestling stage, we conducted a two-by-two factorial experiment by manipulating parasite presence and nest temperature (Ingala et al. 2022). For our study, we followed 14 parasitized-heated and 14 parasitized-non-heated bluebird nests and 12 non-parasitized-heated and 12 non-parasitized non-heated bluebird nests. We also followed nine parasitized-heated and nine parasitized-non heated swallow nests and nine non-parasitized heated and eight non-parasitized non-heated swallow nests.

The experiment was initiated when at least one hatchling was found in the nest and determined to be 0 (hatch age) or 1 day old. For the control treatment, nest material was treated with sterile water to allow for natural parasitism (parasitized). For the experimental treatment, nest material was treated with a 1% permethrin solution (Permectrin II) to remove all parasites (non-parasitized) (Knutie et al. 2016, DeSimone et al. 2018). Nest boxes were revisited two days later to treat nests again to ensure that the treatments were effective. Previous studies found sub-lethal effects of permethrin on nestlings when they were in direct contact with the insecticide (Bulgarella et al. 2020). Therefore, we briefly removed nestlings and the top liner of the nest during treatment application so that nestlings never came into direct contact with permethrin. Parasite treatment was initially determined by a coin flip separately for each species, and all subsequent nests for each species were then alternately assigned to a treatment.

For the heat treatment, we used a metal spatula to lift nest material from the bottom of the box and placed a fresh UniHeat 72+ Hour heat pack (heated) or an exhausted heat pack (non-heated) in the open space (Fig. S1). The packs contained a mixture of charcoal, iron powder, vermiculite, salt, sawdust, and moisture, and based on a past study, elevated temperatures by ±5°C compared to non-heated nests for two days when exposed to the air (Dawson et al. 2005). Nest boxes were revisited every two days to replace active heat packs so that nest boxes continually had elevated temperatures while nestlings were 0 to 10 days old, or to lift nest material with a metal spatula to cause similar disturbance in non-heated, control nests. Heat packs were always checked for parasites before they were removed; any parasites that were found on the heat pack were returned to the nest. Heat treatment was initially determined by a coin flip separately for each species and all subsequent nests were then alternately assigned to a treatment. To record internal nest temperature a data logger (Thermochron iButton DS1921G, Dallas Semiconductor, USA) was placed under the nest liner. Data loggers were programmed to record internal nest temperature once every hour from day of treatment until nests were collected after the nest was empty. We collected sufficient temperature data from 17 bluebird nests but excluded data from swallow nests because only three nests had sufficient data.

### 2.3 Nestling growth metrics and survival

At hatching (0-1 days old), nestlings were weighed to the nearest 0.1 g using a portable digital scale balance (Ohaus CL2000). Nests were revisited when nestlings were between approximately 9-10 days old to weigh nestlings and to measure their tarsus length (to the 0.01 mm), bill length (0.01 mm), and first primary feather length (0.01 mm) using analog dial calipers from Avinet. During this visit, nestlings were also banded with a uniquely numbered USGS metal band (Master’s banding #23623 [SAK]). Beginning when nestlings were approximately 15 days old, the boxes were checked every other day from a distance using binoculars (to prevent pre-mature fledging) and when the nest box was empty, the age at fledging was recorded.

### 2.4 Blood collection and hemoglobin levels

When nestlings were 9-10 days old, a small blood sample (∼20µL) was collected from the brachial vein of each nestling using a 30-gauge sterile needle. Hemoglobin levels (g/dl) of a single individual in each nest were quantified from the whole blood using a HemoCue® HB +201 portable analyzer.

### 2.5 Quantifying parasite abundance

When the nest was empty, we carefully removed nests, expired heat packs, and iButtons from the nest boxes and stored contents in a gallon-sized, labeled plastic bag. Nest material was dissected over trays lined with white paper. All *P. sialia* larvae (1^st^, 2^nd^, and 3^rd^ instars), pupae, and pupal cases were counted to determine a combined total parasite abundance for each nest. After dissection, heat packs were discarded and nest material was placed back into the labeled plastic bags, then weighed on a digital scale to determine the dry nest mass (g) for each nest. Larvae and pupae found in each nest were reared to adulthood in the lab to confirm identification. Macroparasite size can be used as a measurement of virulence (Vale et al. 2011) thus, we calculated pupal volume as a measurement of size. Briefly, the length and width (0.01 mm) of up to ten pupae per nest were haphazardly selected and measured with digital calipers. These measurements were used to calculate pupal volume (V = π*[0.5*width]^2^*length).

### 2.6 Statistical Analyses

To confirm that heat treatment effectively increased temperatures in bluebird nests, we modeled nest temperature over 24 hours in heated and non-heated nests using a general additive mixed model (GAMM) with the gamm function in the ‘*mgcv*’ package (Wood 2011) in R (2019, version 3.6.1.). In this model, the predictors included heat treatment and smoothed term of time in hours and the response variable was hourly nest temperature The effect of time on nest temperature was allowed to vary according to heat treatment. We accounted for the non-independence of temperature observations within nests and Julian hatch dates by including random intercept terms for nest and Julian hatch date. We included data only from time of nest manipulation to fledging for analysis.

To determine the effect of temperature treatment on parasite metrics we quantified parasite abundance density for each nest. Parasite abundance was quantified as the total number of larvae, pupae, and pupal cases counted within each nest. Parasite density was quantified as the number of parasites divided by the total mass of nestlings within a nest (Grab et al. 2019). We calculated total mass for each nest by multiplying the average clutch size for each population by the average hatch mass of swallows (2.4g) and bluebirds (3.4g); average hatch mass was calculated from previous years (2016-2017) at our Minnesota field site. To evaluate the effect of heat treatment on parasite abundance, density, and volume across bird species, we used a generalized linear model (GLM) with a Poisson error distribution for abundance and linear models (LM) with Gaussian distributions for density and volume. The response variables included parasite abundance and density, and the predictors were heat treatment, host species, and the interaction between heat treatment and host species. These models included two covariates, which were log-transformed nest mass and Julian hatch date and excluded non-parasitized nests since no parasites were found in the nests. The parasite volume model also included parasite abundance as a covariate, to account for the intraspecific competition among parasites which might also affect size.

To examine the effects of parasite and heat treatments on fledging success across host species, we used two separate logistic regressions, which included one model for each species. In each model, the response variable was proportion of nestlings that successfully fledged (number of nestlings that survived to the end of the observation period, number of nestlings that died), and the predictors were heat treatment, parasite treatment, and the interaction between heat and parasite treatments. These models included two covariates, which were log-transformed nest mass and Julian hatch date.

To test for the effects of heat and parasite treatments on nestling morphometrics, we used a permutational analysis of variance (PERMANOVA) model for each species because these metrics are non-independent. For each model, the response variable was a resemblance matrix constructed with Euclidean distances of normalized values of average nestling mass, bill length, tarsus length, and first primary feather length of hosts within nests, and the predictors were heat treatment, parasite treatment, and the interaction between heat and parasite treatments. Both models included two covariates, which were log-transformed nest mass and Julian hatch date.

We evaluated 9999 permutations using residuals under a reduced model and examined test statistics associated with Type I sums of squares to determine if there were any effect of treatments after accounting for the effects of the two covariates. To visualize the results of the PERMANOVA, we used distance-based redundancy analysis (dbRDA). The dbRDA was based on an appropriate resemblance matrix as previously described. The underlying predictors were parasite and heat treatments (as categorical variables) and log-transformed nest mass and Julian hatch date (as continuous variables). In the dbRDA plot, we show the centroid values for the four experimental treatments. Ellipses surrounding points represent 95% confidence intervals of groups based on standard errors and were made using the ordiellipse function in the ‘*vegan*’ package (Oksanen et al. 2020) in R. PERMANOVA models and the dbRDA were completed using PERMANOVA+ for PRIMER version 7 (PRIMER-E Ltd, Plymouth, UK). For ease of visualization of the dbRDA point and vector plots, data from PERMANOVA+ for PRIMER were exported, and plots were made using ‘*ggplot2*’ package (Wickham 2016) in R.

Linear mixed-effects models (i.e., univariate tests) with Gaussian distributions were then used to determine the effect of heat treatment, parasite treatment, and their interaction (predictors) on body mass, bill length, tarsus length, and first primary feather length (response variables). All univariate models also included two covariates, which were log-transformed nest mass and Julian hatch date, and a random intercept term of nest ID.

To test for the effects of heat and parasite treatments on nestling hemoglobin, we used linear models with Gaussian distributions for each species. For each model the response variable was nestling hemoglobin level (only one nestling per nest) and the predictors were heat treatment, parasite treatment, and the interaction between heat and parasite treatments. In addition, in each model body mass was a covariate to control for the difference in size of bird species. To examine how parasite abundance contributed to these patterns, we used two additional linear models with Gaussian distributions for each species. Models included hemoglobin level as the response variable and heat treatment, parasite treatment, and their interaction as the predictors. For all univariate models, test statistics associated with Type III sums of squares were evaluated.

## 3 RESULTS

### 3.1 Effect of temperature treatment on parasite metrics

For bluebirds, heated nests had higher nest temperatures compared to nests that were non-heated (*t* = -2.863, *P* = 0.004, Fig. 2). For both species, treating nests with permethrin was effective at eliminating parasites; no parasites were found in any nests treated with permethrin. Heat treatment, host species, and the interaction between heat treatment and host species affected parasite abundance (Table S1). For bluebirds, parasite abundance was higher in heated nests compared to non-heated nests (Fig 3A). For swallows, parasite abundance was lower in heated nests compared to non-heated nests (Fig 3B). Host species and the interaction between heat treatment and host species, but not heat treatment alone, affected parasite density (Table S1). For bluebirds, parasite density was higher in heated nests compared to non-heated nests (Fig 3C). For swallows, parasite density was lower in heated nests compared to non-heated nests (Fig 3D). Heat treatment, host species, and the interaction between heat treatment and host species did not affect blowfly pupal size (Table S1).

**Fig 2.**
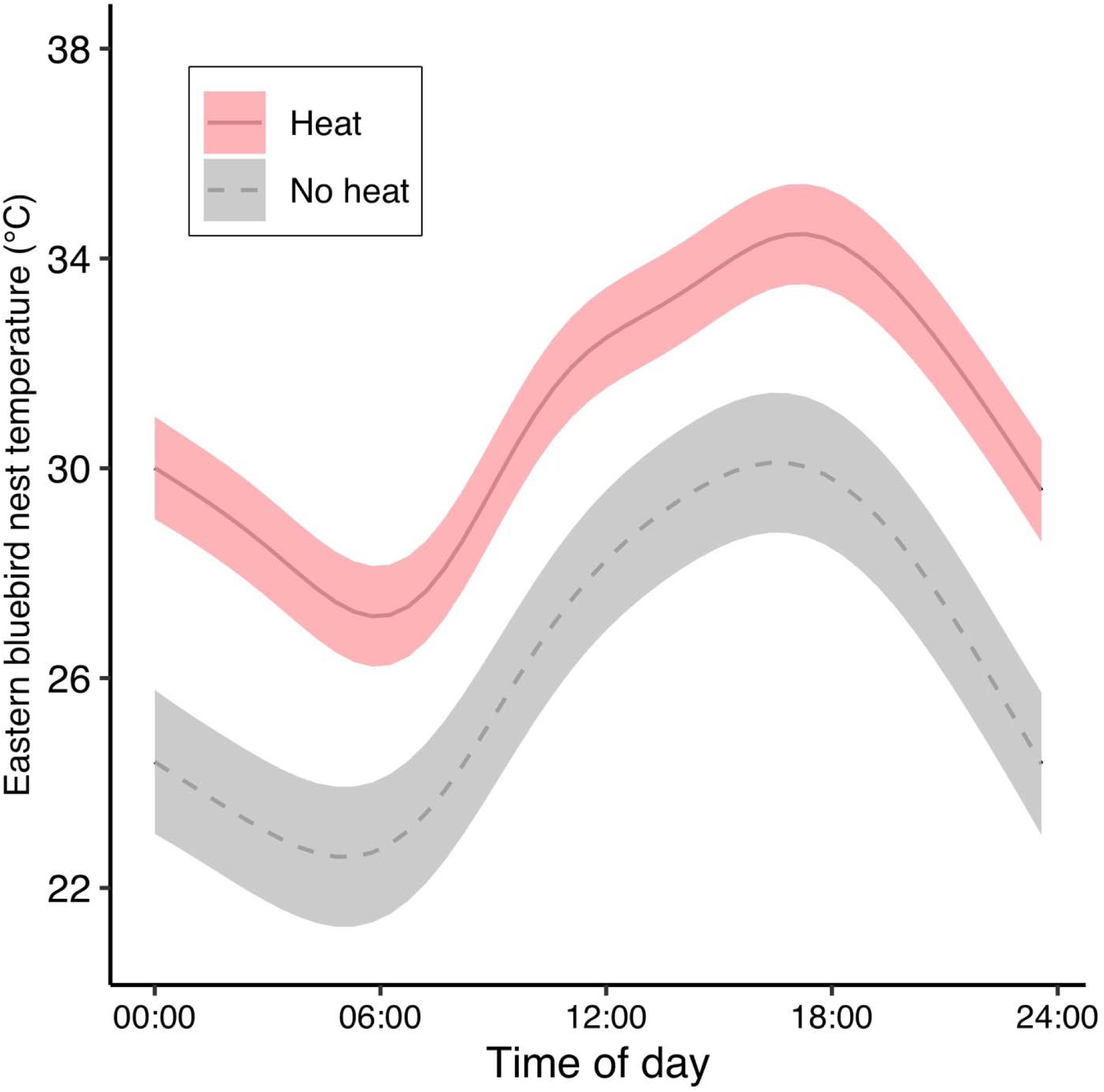
The effect of heat treatment on nest temperature. In bluebird nests, heat increased nest temperatures throughout the course of a day. Lines are predictions from generalized additive mixed models, which included random intercept terms for nest and Julian hatch date to account for the non-independence of individual observations within nests and Julian hatch dates. Error ribbons are standard errors of model predictions.

**Fig 3.**
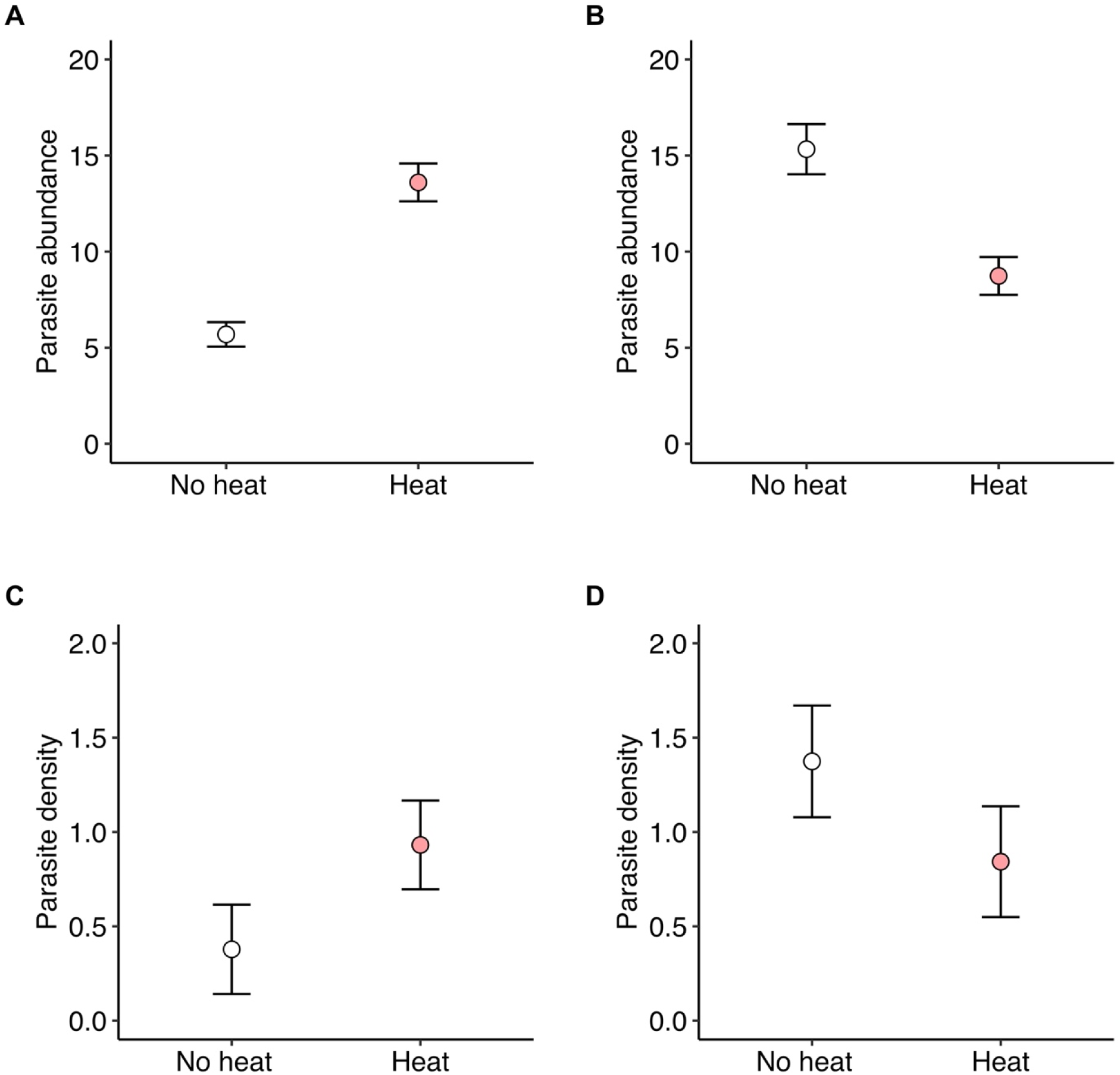
Heat treatment on parasite fitness demonstrates a different directionality of parasite fitness across the two host species. A) Parasite abundance increased in bluebird nests and B) decreased in tree swallow nests in response to heat treatment. C) Parasite density (total parasites/gram of nestling) increased in bluebird nests and D) decreased in tree swallow nests in response to heat treatment. Standard errors shown.

### 3.2 Fledging success and nestling growth

Heat treatment, parasite treatment, and their interaction did not affect bluebird survival (Fig. 4A, Table S2). The interaction between heat and parasite treatments, but not the main effects of heat or parasite treatments, affected swallow survival (Fig. 4B, Table S2). Overall, swallow survival was higher with heated nests compared to non-heated nests across both parasite treatments.

**Fig. 4.**
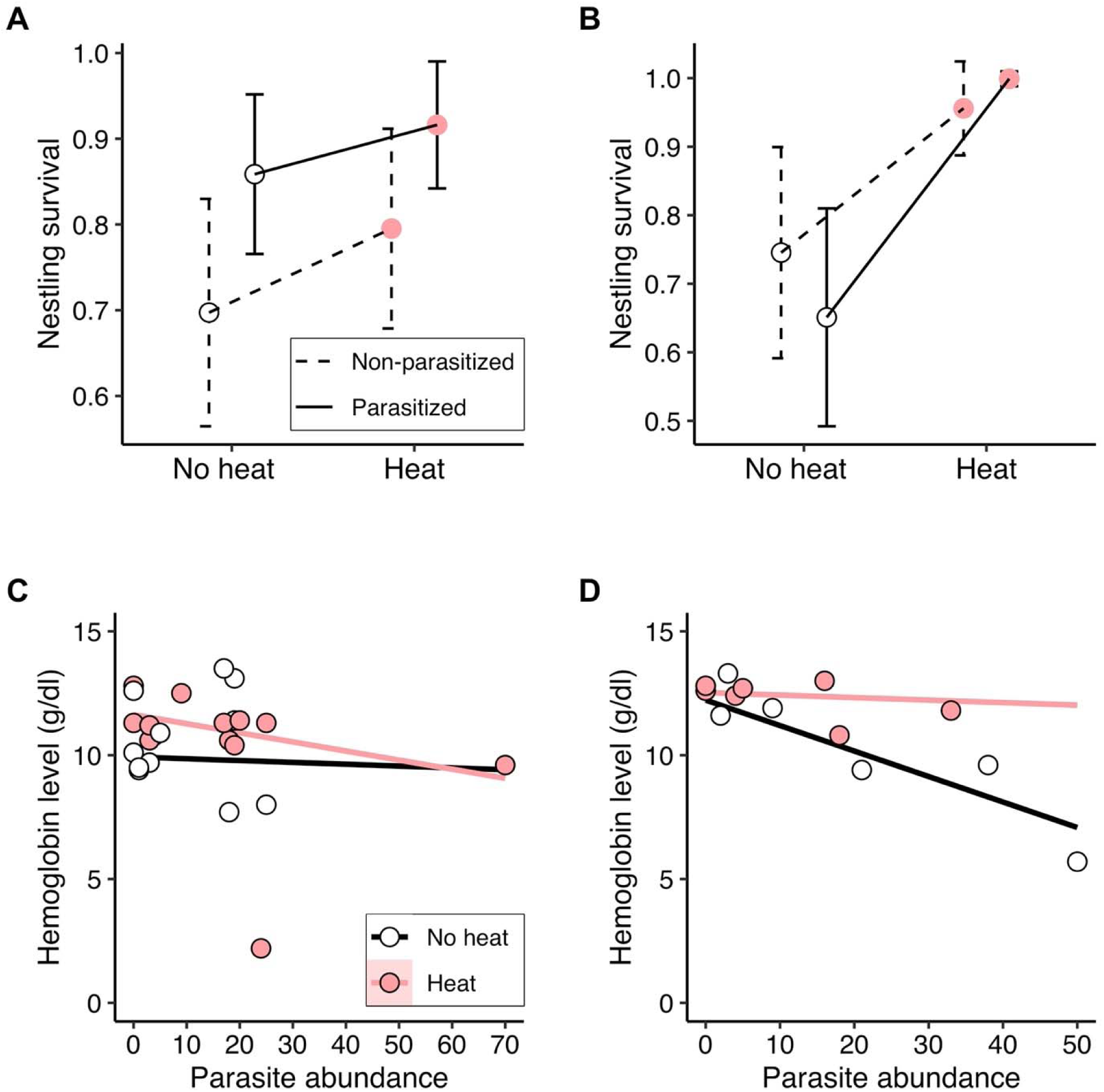
The effect of heat treatment and parasitism on nestling survival and blood loss. A) Heat treatment, parasitism, and their interaction affected bluebird survival. B) However, swallow survival was higher in heated nests compared to non-heated nests. Means and binomial standard errors are shown. C) Heat treatment, parasitism, and their interaction affected bluebird hemoglobin levels. D) Without heat, tree swallow hemoglobin levels declined as parasite abundance increased. With heat, hemoglobin levels did not decline despite increasing parasite abundance.

Heat treatment, parasite treatment, and their interaction did not affect swallow morphometrics (Table S3, S4). Bluebird morphometrics were driven by a marginal effect of heat and the interaction between parasite and heat treatments, but not the main effect of parasite treatment (Table S3, S4). Heat treatment had the most significant effect on bluebird morphometrics (Fig. S3A). Bluebirds from non-heated nests had larger body masses compared to nestlings from heated nests (Fig. S2). Based on the univariate tests, heat treatment, parasite treatment, and their interaction did not affect bill length or tarsus length (Table S3, Table S5). However, the interaction between parasite treatment and heat treatment, but not their main effects, affected primary feather length (Table S3, Table S5). Within non-parasitized nests, nestlings from non-heated nests had significantly longer first primary feathers compared to nestlings from heated nests, and this effect was reversed when nests were parasitized.

### 3.3 Nestling hemoglobin

Heat treatment, parasite treatment, and their interaction did not affect bluebird hemoglobin levels (Table S3, Table S6). Furthermore, parasite abundance did not affect hemoglobin levels across heat treatments. (Fig. 4C, Table S7). For swallows, there was a marginal effect of parasite treatment on hemoglobin levels, but no effect of heat treatment or their interaction (Table S6). Hemoglobin levels on average were lower for swallows in parasitized nests compared to non-parasitized nests. Importantly, parasite abundance and the interaction between parasite abundance and heat treatment did effect swallow hemoglobin levels (Fig. 4D, Table S7). For swallow nestlings from non-heated nests, hemoglobin levels declined as parasite abundance increased. However, for nestlings from heated nests, hemoglobin levels remained stable as parasite abundance increased (Fig. 4D).

## 4 DISCUSSION

Our study demonstrates heat can have varying effects on host-parasite interactions depending on host species. When heated, bluebird nests had more parasites and swallow nests had fewer parasites, compared to non-heated nests. Interestingly, neither heat nor parasite abundance affected hemoglobin levels in bluebirds, while swallows from heated nests were more tolerant to blood loss in response to increasing parasite abundance, compared to non-heated nests. Our results show that there are direct effects of elevated temperature on hosts in their response to parasitism, but these effects vary across species.

We found that the effect of heat on differing parasite abundance across hosts was likely mediated by an indirect effect through host defense strategies. For example, heat might have strengthened resistance to parasites via the IgY antibody response in swallows compared to nestlings from non-heated nests (DeSimone et al. 2018, Grab et al. 2019). After an individual is bitten, inflammatory pathways are activated and IgY antibodies are produced, which can bind to larval blowflies (Koop et al. 2013, Owen et al. 2010). Inflammation can negatively affect the blowflies by causing tissue swelling, preventing the parasites from feeding. Surprisingly, non-heated swallow nests had more parasites than non-heated bluebird nests, which contrasts with past patterns in these populations (bluebirds had twice as many parasites per gram of nestling compared to swallows) (Grab et al. 2019). This difference suggests that host defenses against blowflies might vary annually. During previous years, bluebirds from this northern Minnesota population generally did not produce a detectable IgY immune response to the parasites (Grab et al. 2019), unless supplemented with food (Knutie, 2020). Food resource availability could have been higher in 2018 facilitating higher resistance and fewer parasites in non-heated bluebird nests. However, heat could have counteracted this resource benefit by reducing resistance in bluebirds. Other studies have found that heat can depress the activity of the immune system (Calefi et al. 2016). For example, white blood cell count and humoral response were inhibited in heat stressed commercial hens because of a reduction in leukocyte activity and antibody synthesis (Mashaly et al. 2004). In our study, we did not quantify immune metrics in nestlings, which could have helped explain our results. Future studies are still needed to understand the mechanisms to how heat specifically could be influencing host resistance in this system.

Overall, tree swallow nestling survival was higher in heated nests, even when nestlings were parasitized. Other studies have similarly found that tree swallow nestlings can have higher survival in response to elevated temperatures, which support our findings (Dawson, Lawrie, O’Brien, 2005, McCarthy & Winkler, 1999). One potential mechanism for higher survival is that heated nests allowed nestlings to devote less energy toward maintaining an optimal body temperature and more energy toward growth and immunity to parasitism (Ganeshan et al. 2019). In contrast, neither parasitism nor heat influenced bluebird nestling survival. However, we found an interactive effect of parasitism and heat on bluebird growth. Parasitized bluebird nestlings had longer first primary feathers with heat. Similar to our study, Murphy (1985) found eastern kingbirds (*Tyrannus tyrannus*) had longer first primary feathers, but less mass gained, in response to higher ambient temperatures. Other studies have proposed that primary flight feather growth is prioritized in nestling development to facilitate successful or earlier fledging (Saino et al. 1998, Andreasson et al. 2017). When dealing with parasitism, the growth of first primary feathers in bluebirds may be prioritized to help fledge sooner and therefore escape parasitism, and this growth could be accommodated by elevated temperatures.

Swallow nestlings from heated nests maintained high hemoglobin levels (lower blood loss) despite increasing parasite abundances. This result suggests that swallows are tolerant of the sublethal effects of parasitism when exposed to elevated temperatures. Previous studies have found that parasitism can substantially decrease hemoglobin levels, since ectoparasites remove blood from their hosts (Grab et al. 2019, Sun et al. 2019, DeSimone et al. 2018). Without parasites, higher nest temperature can even result in higher hematocrit levels (more red blood cells) in nestling tree swallows (Ardia 2013). Elevated nest temperatures could result in faster red blood cell recovery in nestlings via changing oxygen availability, which could explain our results (Niedojadlo et al. 2018, Fair et al. 2007, Bradley et al. 2020). These results provide support for the emergence of tolerance to parasitism with elevated temperatures.

In this study, we found important implications for the effects of elevated temperature interacting with an effect of parasitism. Specifically, infestation with the same parasite, *P. sialia*, decreased in tree swallow nests and increased in eastern bluebird nests in response to elevated nest temperature. In natural host-parasite interactions, elevated temperatures could have different consequences for the health of host species and potentially alter the balance between defense strategies that the hosts use against infection, thereby changing the dynamics of the relationship with the parasite. Elevated temperatures may also directly affect the parasite apart from, or at the same time as, directly affecting the host. The relationship between temperature and parasitism throughout the interaction presented in this study provides new insights on the role of a changing climate in the future of host-parasite interactions. Differences in host responses may alter the dynamics of each individual interaction, thereby having implications for the overall multi-host-parasite system.

## ACKNOWLEDGMENTS

We thank Steve Knutie and Doug Thompson for building nest boxes and the University of Minnesota Itasca Biological Station and Laboratories for logistical support. We also thank the following people for the interest in our work, along with access to nest boxes located on their property: Lesley Knoll and Aaron Hebbeler, Helen Perry, Doug and Dawn Thompson, Pioneer Farms, and Rock Creek General Store. A special thanks to the members of the Hall lab at Indiana University who contributed generous suggestions toward improving the manuscript.

The authors and collaborators also wish to acknowledge the Ojibwe people who have cared for and occupy the land in which our research was conducted. The University of Minnesota Itasca Biological Station and Laboratories is located on the land ceded by the Mississippi and Pillager Bands of Ojibwe in the Treaty of Washington, commonly known as the 1855 Treaty. This treaty affirms the reserved rights doctrine and the inalienable rights of Ojibwe people to uphold their interminable relationship to the land. With affiliation to the aforementioned academic institutions, it is our responsibility to acknowledge Native rights and the institutions’ history with them. We are committed to continue building relationships with the Ojibwe People through recognition, support, and to advocate for all Native American Nations. We strive to be good stewards of our place and privilege. *This land acknowledgement was revised and written with support from Rebecca Dallinger and Joe Allen. It is a living document open to changes*.

## DECLARATIONS

### ETHICS APPROVAL

All applicable institutional guidelines for the care and use of animals were followed (University of Connecticut IACUC protocol #A18-005).

### CONSENT TO PARTICIPATE

Not applicable

### CONSENT FOR PUBLICATION

Not applicable

### DATA AVAILABILITY STATEMENT

Data will be made available via FigShare upon acceptance.

### CODE AVAILABILITY

Code will be made available via FigShare upon acceptance.

### AUTHOR CONTRIBUTIONS

LA and SAK conceived the ideas and designed methodology; LA, AP, SAK collected data; SR and GV analyzed the data; LA, SAK, GV, SR contributed to the writing of the manuscript. All authors gave final approval of the manuscript.

