## Supplemental Materials for "Elevated nest temperature has opposing effects on host species infested with parasitic nest flies"

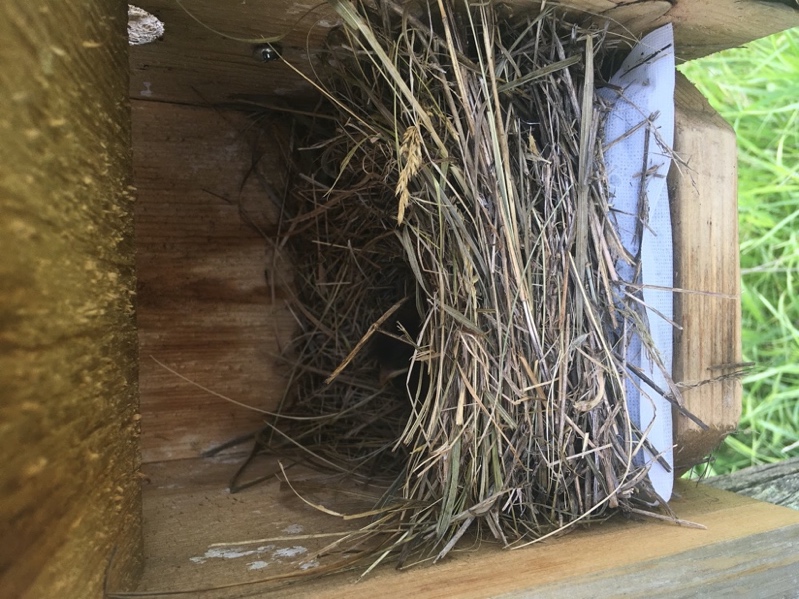
 Figure S1. Experimental heat pack inserted under eastern bluebird nest in nest box.

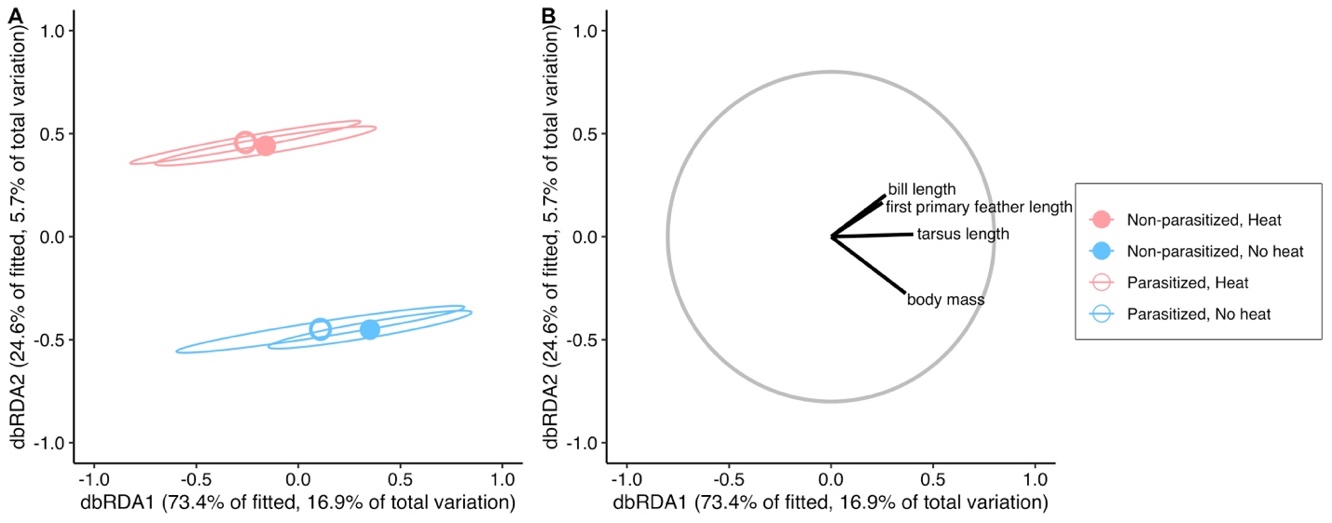

Figure S2. The effect of heat and parasite treatments on multivariate size and mass of eastern bluebirds. A) Distance-based redundancy analysis (dbRDA) plot of multivariate eastern bluebird size and mass showing differences between heat treatments. Eastern bluebirds tend to weigh less in nests that were heated. Individual points correspond to the four treatments in the experiment. Ellipses are 95% confidence intervals. B) Vector overlay of size and mass responses for the corresponding dbRDA plot. The gray circle corresponds to vector lengths that would have a correlation coefficient of one with each axis.

Table S1. Effect of heat treatment, bird species, and the interaction on parasite abundance, density, and size.

| Variables | *df* | Test Statistic | *P* |
| --- | --- | --- | --- |
| **Parasite abundance** |  | Wald *χ*^2^ |  |
| log(nest mass) | 1 | 86.978 | **<0.001** |
| Day first egg hatched | 1 | 43.117 | **<0.001** |
| Heat treatment | 1 | 52.081 | **<0.001** |
| Bird species | 1 | 56.739 | **<0.001** |
| Heat treatment x bird species | 1 | 62.024 | **<0.001** |
| Error | 40 |  |  |
| **Parasite density** |  | *F* |  |
| log(nest mass) | 1 | 9.057 | **0.005** |
| Day first egg hatched | 1 | 5.380 | **0.026** |
| Heat treatment | 1 | 2.785 | 0.103 |
| Bird species | 1 | 6.656 | **0.014** |
| Heat treatment x bird species | 1 | 4.229 | **0.046** |
| Error | 40 |  |  |
| **Parasite size** |  | Wald *F* |  |
| Parasite abundance | 1, 14.772 | 0.715 | 0.411 |
| Heat treatment | 1, 15.112 | 0.347 | 0.564 |
| Bird species | 1, 16.617 | 0.355 | 0.560 |
| Heat treatment x bird species | 1, 15.761 | 0.031 | 0.863 |

Table S2. Effect of heat treatment, parasite treatment, and the interaction on nestling survival.

| Variables | *df* | Wald *χ*^2^ | *P* |
| --- | --- | --- | --- |
| **Eastern bluebird** |  |  |  |
| log(nest mass) | 1 | 26.951 | <0.001 |
| Day first egg hatched | 1 | 4.367 | 0.037 |
| Heat treatment | 1 | 0.982 | 0.322 |
| Parasite treatment | 1 | 3.523 | 0.061 |
| Parasite treatment x heat treatment | 1 | 0.008 | 0.931 |
| Error | 45 |  |  |
| **Tree Swallow** |  |  |  |
| log(nest mass) | 1 | 20.286 | <0.001 |
| Day first egg hatched | 1 | 3.257 | 0.071 |
| Heat treatment | 1 | 3.293 | 0.070 |
| Parasite treatment | 1 | 0.414 | 0.520 |
| Parasite treatment x heat treatment | 1 | 6.979 | 0.008 |
| Error | 29 |  |  |

Table S3. Effect of parasite and heat treatment on bluebird and swallow nestling mass, size, and physiology. Numbers are mean ± SE and numbers in parentheses are numbers of nests.

|  | Parasitized | |  | Non-parasitized | |
| --- | --- | --- | --- | --- | --- |
| Measurement | No Heat | Heat |  | No Heat | Heat |
| **Eastern Bluebird** |  |  |  |  |  |
| Tarsus (mm) | 18.62 ± 0.30 (12) | 18.70 ± 0.17 (13) |  | 19.11 ± 0.19 (10) | 18.56 ± 0.29 (10) |
| 1^st^ Primary (mm) | 13.94 ± 1.09 (12) | 17.12 ± 0.88 (13) |  | 16.58 ± 0.78 (10) | 14.51 ± 0.99 (10) |
| Bill length (mm) | 5.17 ± 0.11 (12) | 5.39 ± 0.08 (13) |  | 5.34 ± 0.07 (10) | 5.24 ± 0.09 (10) |
| Mass (g) | 26.11 ± 0.75 (12) | 24.21 ± 0.66 (13) |  | 26.65 ± 0.72 (10) | 23.97 ± 0.56 (10) |
| Hemoglobin (g/dl) | 10.54 ± 0.59 (11) | 10.46 ± 0.73 (13) |  | 11.78 ± 0.49 (10) | 10.86 ± 0.22 (9) |
| **Tree Swallow** |  |  |  |  |  |
| Tarsus (mm) | 11.59 ± 0.17 (7) | 11.57 ± 0.14 (8) |  | 11.60 ± 0.08 (8) | 11.55 ± 0.19 (7) |
| 1^st^ Primary (mm) | 12.34 ± 1.02 (7) | 13.66 ± 1.30 (8) |  | 12.88 ± 1.41 (8) | 15.40 ± 1.05 (7) |
| Bill length (mm) | 4.29 ± 0.12 (7) | 4.53 ± 0.07 (8) |  | 4.45 ± 0.14 (8) | 4.70 ± 0.17 (7) |
| Mass (g) | 19.45 ± 0.69 (7) | 19.73 ± 0.74 (8) |  | 20.76 ± 0.85 (8) | 20.89 ± 0.55 (7) |
| Hemoglobin (g/dl) | 10.25 ± 1.09 (6) | 12.30 ± 0.29 (7) |  | 11.74 ± 0.37 (8) | 12.32 ± 0.18 (6) |

Table S4. Nestling multivariate growth analysis.

| Variables | *df* | Test Statistic | *P* |
| --- | --- | --- | --- |
| **Eastern bluebird** |  | Pseudo *F* |  |
| log(nest mass) | 1 | 2.135 | 0.121 |
| Day first egg hatched | 1 | 7.211 | 0.002 |
| Heat treatment | 1 | 2.837 | 0.057 |
| Parasite treatment | 1 | 0.101 | 0.959 |
| Parasite treatment x heat treatment | 1 | 3.165 | 0.049 |
| Error | 38 |  |  |
| **Tree swallow** |  | Pseudo *F* |  |
| log(nest mass) | 1 | 0.457 | 0.678 |
| Day first egg hatched | 1 | 2.287 | 0.098 |
| Heat treatment | 1 | 1.980 | 0.131 |
| Parasite treatment | 1 | 1.478 | 0.231 |
| Parasite treatment x heat treatment | 1 | 0.060 | 0.984 |
| Error | 24 |  |  |

Table S5. Eastern bluebird univariate growth analysis.

| Variables | *df_num_*, *df_den_* | Wald *F* | *P* |
| --- | --- | --- | --- |
| **Bill length** |  |  |  |
| log(nest mass) | 1, 38.696 | 0.133 | 0.717 |
| Day first egg hatched | 1, 39.181 | 6.848 | 0.013 |
| Heat treatment | 1, 38.209 | 0.143 | 0.707 |
| Parasite treatment | 1, 37.176 | 1.275 | 0.266 |
| Parasite treatment x heat treatment | 1, 38.176 | 2.575 | 0.117 |
| **Tarsus length** |  |  |  |
| log(nest mass) | 1, 38.852 | 2.504 | 0.122 |
| Day first egg hatched | 1, 39.79 | 7.564 | 0.009 |
| Heat treatment | 1, 38.198 | 1.372 | 0.249 |
| Parasite treatment | 1, 36.792 | 1.530 | 0.224 |
| Parasite treatment x heat treatment | 1, 38.175 | 1.540 | 0.222 |
| **First primary length** |  |  |  |
| log(nest mass) | 1, 38.628 | 1.628 | 0.210 |
| Day first egg hatched | 1, 39.005 | 3.780 | 0.059 |
| Heat treatment | 1, 38.199 | 1.481 | 0.231 |
| Parasite treatment | 1, 37.291 | 3.345 | 0.075 |
| Parasite treatment x heat treatment | 1, 38.165 | 7.744 | 0.008 |
| **Mass** |  |  |  |
| log(nest mass) | 1, 38.704 | 1.378 | 0.248 |
| Day first egg hatched | 1, 39.205 | 4.389 | 0.043 |
| Heat treatment | 1, 38.210 | 5.453 | 0.025 |
| Parasite treatment | 1, 37.161 | 0.174 | 0.679 |
| Parasite treatment x heat treatment | 1, 38.177 | 0.302 | 0.586 |

Table S6. Effect of heat treatment, parasite treatment, and the interaction of nestling hemoglobin.

| Variables | *df* | *F* | *P* |
| --- | --- | --- | --- |
| **Eastern bluebird** |  |  |  |
| Body mass | 1 | 8.041 | 0.007 |
| Heat treatment | 1 | 0.317 | 0.577 |
| Parasite treatment | 1 | 1.703 | 0.200 |
| Parasite treatment x heat treatment | 1 | 0.062 | 0.805 |
| Error | 38 |  |  |
| **Tree swallow** |  |  |  |
| Body mass | 1 | 6.028 | 0.022 |
| Heat treatment | 1 | 0.233 | 0.634 |
| Parasite treatment | 1 | 4.170 | 0.053 |
| Parasite treatment x heat treatment | 1 | 3.450 | 0.077 |
| Error | 22 |  |  |

Table S7. Effect of heat treatment, parasite abundance, and the interaction on nestling hemoglobin.

| Variables | *df* | *F* | *P* |
| --- | --- | --- | --- |
| **Eastern bluebird** |  |  |  |
| Body mass | 1 | 5.439 | 0.031 |
| Heat treatment | 1 | 1.654 | 0.214 |
| Parasite abundance | 1 | 0.011 | 0.917 |
| Parasite abundance x heat treatment | 1 | 0.142 | 0.711 |
| Error | 19 |  |  |
| **Tree swallow** |  |  |  |
| Body mass | 1 | 5.317 | 0.050 |
| Heat treatment | 1 | 0.216 | 0.655 |
| Parasite abundance | 1 | 27.092 | 0.001 |
| Parasite abundance x heat treatment | 1 | 8.791 | 0.018 |
| Error | 8 |  |  |
